# Embedded control of cell growth using tunable genetic systems

**DOI:** 10.1101/2021.06.28.450134

**Authors:** Virginia Fusco, Davide Salzano, Davide Fiore, Mario di Bernardo

**Author notes:** **Correspondence** Mario di Bernardo.

## Abstract

We present an embedded feedback control strategy to control the density of a bacterial population, allowing cells to self-regulate their growth rate so as to reach a desired density at steady state. We consider a static culture condition, where cells are provided with a limited amount of space and nutrients. The control strategy is built using a tunable expression system (TES), which controls the production of a growth inhibitor protein, complemented with a quorum sensing mechanism for the sensing of the population density. We show on a simplified population-level model that the TES endows the control system with additional flexibility by allowing the set-point to be changed online. Finally, we validate the effectiveness of the proposed control strategy by means of realistic *in silico* experiments conducted in BSim, an agent-based simulator explicitly designed to simulate bacterial populations, and we test the robustness of our design to disturbances and parameters’ variations due, for instance, to cell-to-cell variability.

## 1 INTRODUCTION

Microorganisms, such as bacteria and yeast, are widely used in biotechnology to produce proteins and chemicals of industrial and biomedical interests ^1–6^. This is made possible by embedding artificial genetic circuits into cells modifying their natural behavior, that is, by engineering *ad hoc* at what conditions and rates the genes of interest are expressed into the cells. In this context a key open problem is to channel cell resources from biomass production (i.e., cell growth and division) to protein production while preventing accumulation of toxic by-products ^7–11^. Specifically, the cell growth rate needs to be carefully regulated in order to make the biomass density in the bioreactors reach and maintain a desired reference value so as to guarantee higher efficiency of the bioproduction process. This objective is generally achieved by means of external systems controlling the density of the cell populations inside the bioreactors ^12–17^. However, designing cellular populations capable of self-regulating their growth rate by means of embedded genetic feedback mechanisms could provide intrinsic robustness to the bioproduction process, e.g., preventing extinction or starvation, while simplifying the design of external control systems.

A pioneering solution for the self-regulation of population density in single strain microbial populations was proposed in 18 where a quorum sensing based mechanism is used to self-limit the number of the cells. Alternative approaches involve engineering multiple microbial populations to achieve self-regulation of their relative numbers ^19–24^ and in some cases also regulation of both the consortium size and composition ^25,26^. In all previous work, the desired population densities are hard-wired into the design of the synthetic gene regulatory networks. Hence, achieving a different working condition requires re-engineering the entire consortium.

In this paper, inspired by 18, we present a genetic feedback strategy for population control. Specifically, by means of a quorum sensing mechanism, which allows cells to sense the population density, we render cells capable of self-regulating their own growth rate by producing a growth inhibitor protein. The genetic controller governing this process inside the cells is implemented by means of a *tunable expression system* (TES), recently proposed in 27. By exploiting this circuit, the production rate of the inhibitor protein can be dynamically changed depending on the concentration of the quorum sensing molecule present into the environment. In so doing, a feedback mechanism can be embedded into the cells so that they become able to regulate their own relative numbers. Thanks to its characteristic features, the TES provides additional flexibility to the control system by allowing the desired set-point for the population density to be changed online according to some exogenous input, and to compensate for unavoidable inaccuracies in the offline tuning of the system parameters, and uncertainties due to cell-to-cell variability.

After presenting the details of the proposed genetic controller, we analyze its steady-state response by considering a simplified model obtained by averaging the dynamics over the entire population assumed to grow in a culture tube. This allows us to analytically derive an approximate relationship between some tunable parameter and the steady-state population density that we then exploit for control design. Finally, the performance and the robustness of the proposed control system are validated *in silico* by means of realistic agent-based simulations performed in BSim ^28,29^, a platform to simulate microbial populations which was originally designed for microfluidic experiments only and is adapted here to simulate cells growing in test tube cultures.

## 2 POPULATION CONTROL

The control strategy we present here consists of closing a feedback loop between the cell population and its environment allowing the population to self-regulate its own density by producing a growth inhibitor protein (Figure 1).

**FIGURE 1.**
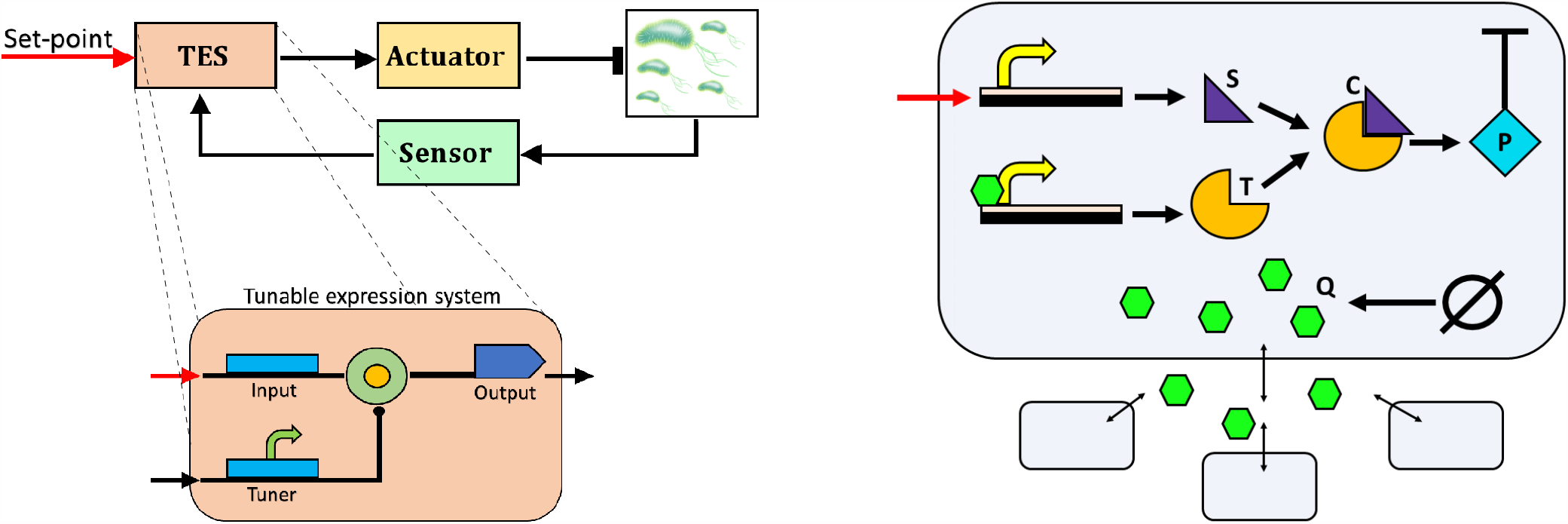
Genetic feedback control of the cell population density. *Left panel:* Representation of the population control strategy. The cell population can self-regulate its growth rate by sensing its current density in the culture environment and comparing it to a desired set-point. This comparison is realized by means of a tunable expression system (TES), which allows its input-output relationship to be dynamically changed as a function of the tuner signal, so that its output can modulate the production of a growth inhibitor protein, closing the feedback loop. *Right panel:* Abstract biological implementation scheme of the TES. Each cell produces a quorum sensing molecule Q (green hexagons) that diffuses into the environment and in the other cells. The molecule Q, proportional to the population density, regulates the production of the species T, which, reacting with the species S, acts as the tuner signal for the production of the complex C. The production of the growth inhibitor protein P is activated by the complex C, thus closing the loop. Notice that the rate of production of the species S can be either tuned offline or regulated online to specify the desired set-point for the population density *N*_d_.

This is realized by comparing the concentration of a *quorum sensing* molecule, produced by the cells and diffusing into the environment, to some desired set-point value of the population density. The concentration of such diffusing molecule into the environment is proportional to the density of the population, therefore, regulating its concentration, directly affects the number of cells into the environment.

The production of the growth inhibitor protein is activated by a genetic controller implemented by means of a tunable expression system (TES) recently proposed in 27 (Figure 1, left panel). Specifically, this genetic circuit allows the *input-output* relationship between the set-point value and the production rate of the inhibitor protein to be dynamically changed in response to the concentration of the quorum sensing molecule. In this way, the quorum sensing molecule acts as a *tuner* input affecting the production rate of the inhibitor protein, closing the loop. Indeed, when the concentration of the quorum sensing molecule into the environment increases (corresponding to a growth in population density), the production rate of the inhibitor protein also increases and thus the population density decreases. Vice versa, when the population density decreases, production of the inhibitor protein also decreases and the growth rate of the population increases again (Figure 1, right panel).

Notice that the desired set-point value for the population density can be set either online by means of external inputs, e.g., by changing the concentration of some inducer molecule in the growth medium ^30^ or by applying light stimuli via optogenetics ^31,32^, or offline by adequately tuning the parameters of the genetic controller.

### 2.1 Modeling

We assume that the cells are cultured in an environment of volume *V* with limited availability of nutrients, such as a *test tube*. We label with *i, i* = 1, …, *M*(*t*), each cell in the population, where *M*(*t*) : ℝ_≥0_ ↦ ℕ is the total number of cells in the population at time *t*. The dynamics of the genetic control system embedded in each cell *i* and depicted in Figure 1, right panel, is described by a deterministic model we derived from the laws of mass-action and Michaelis-Menten kinetics as the following set of ordinary differential equations:

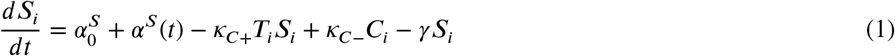

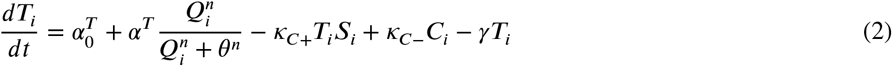

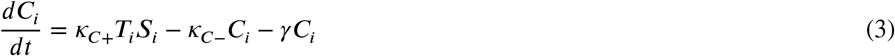

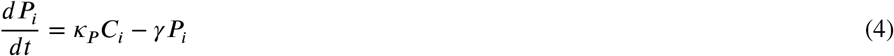

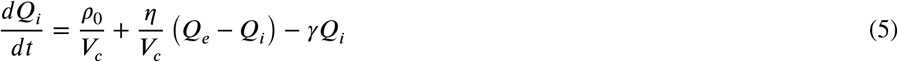

where the state variables *S*_*i*_, *T*_*i*_, *C*_*i*_, *P*_*i*_ denote, respectively, the concentrations inside cell *i* of the toehold switch transcript S, the tuner sRNA T, the switch-sRNA complex C of the TES ^27, SI^, and of the inhibitor protein P. Moreover, *Q*_*i*_ and *Q*_*e*_ denote the concentrations of the quorum sensing molecule Q inside each cell *i* and into the external environment, respectively. That is, *Q*_*i*_ : = *q*_*i*_/*V*_*c*_ and *Q*_*e*_ = *q*_*e*_/*V*_*e*_, where *q*_*i*_ and *q*_*e*_ denote the amount of molecules into each cell with volume *V*_*c*_ and into the environment with volume *V*_*e*_ = *V* − *MV*_*c*_, respectively. For the sake of simplicity, in the above equations we assumed that all cells are identical and all molecular species degrade, mostly because of dilution, following first-order kinetics with rate *γ*.

The dynamics of the TES is described by equations (1)-(3). In (1), the toehold switch S is transcribed with rate 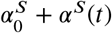, where the value of the function *α*^*S*^ (*t*) encodes the desired set-point value for the population density, which will be assumed constant in the rest of the paper, that is, *α*^*S*^ (*t*) = *α*^*S*^, for all *t >* 0. In (2), the tuner sRNA T is transcribed with rate 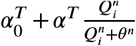, where *θ* is the activation coefficient and is the Hill coefficient, and finally in (3) the switch-sRNA complex C binds and unbinds with rates *κ*_*C*+_ and *κ*_*C*−_, respectively. The inhibitor protein P, with dynamics given in (4), is the output of the TES circuit and is produced at rate *κ*_*P*_ *C*_*i*_. Finally, the quorum sensing molecule Q, whose dynamics is described in (5), is produced by each cell with a constant rate *ρ*_0_, regulating the activation of the transcription of the tuner sRNA T, and diffuses through the cell membrane with diffusion rate *η*, so that the evolution of the quorum sensing molecules into the environment is described by

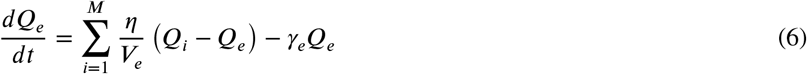

where *γ*_*e*_ is the external degradation rate.

In the following sections, we first present a simpler *aggregate model* of the closed-loop system allowing us to analyze the effects that a change of the reference parameter *α*^*S*^ has on the steady-state value of the population density. Then, we describe the results of an exhaustive set of realistic *in silico* experiments performed by means of the agent-based bacterial simulator BSim ^28,29^ to validate the effectiveness of the proposed approach.

## 3 STEADY-STATE ANALYSIS

Model (1)-(6) can be recast to describe the average behavior of a populations of cells, each embedding the genetic control system described in Section 2. Specifically, denote with *N*(*i*) : ℝ_≥0_ ↦ ℝ_≥0_ the density of the cell population growing in the volume *V* at time *t*, that is, *N*(*t*) = *M*(*t*)/*V*. The *aggregate model* of the cell population dynamics can be derived as (see Appendix A for further details):

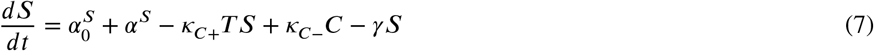

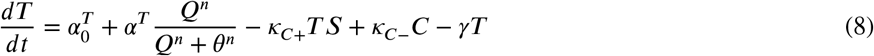

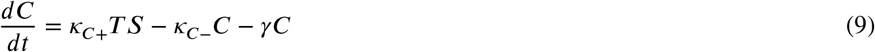

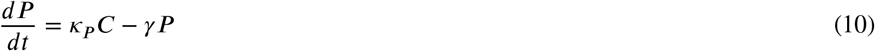

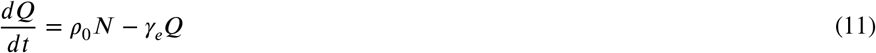

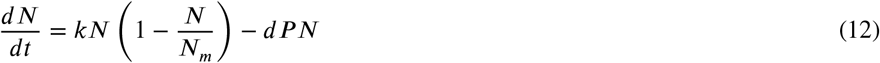

where the state variables *S, T, C, P* denote, respectively, the average concentrations of molecules in each cell of the toehold switch transcript S, the tuner sRNA T, the switch-sRNA complex C of the TES, and of the inhibitor protein P, while *Q* denotes the concentration of quorum sensing molecule Q in the volume *V*, that is, *Q* := *q*/*V*, with *q* being the total amount of Q in the culture volume. The notation and the meaning of the coefficients in (7)-(11) are the same as in (1)-(5). Moreover, in equation (12), *k* denotes the intrinsic growth rate of the population, *N*_*m*_ the carrying capacity, i.e., the maximum density the population can reach given the nutrients available, and *d* the growth inhibition rate due to the protein P.

The aggregate model (7)-(12) allows us to obtain a very simple, yet approximated relationship between the value of the population density, *N*_ss_ = lim_*t* →∞_ *N*(*t*), and the reference parameter *α*^*S*^ at steady state, which is described by the following proposition.

### Proposition 1

Under the assumptions that the basal synthesis rates of the species *S* and *T* are negligible, that is, 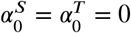, that *n* = 2, and that

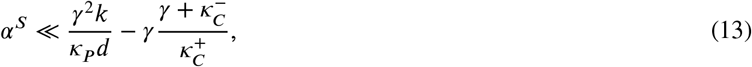

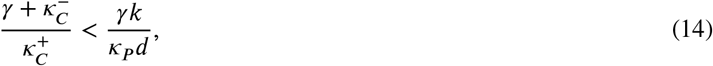

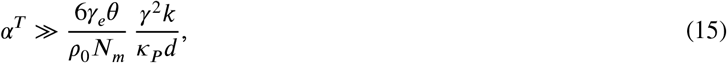

at steady state the value of the population density *N*_ss_, normalized by the carrying capacity *N*_*m*_, can be approximated by

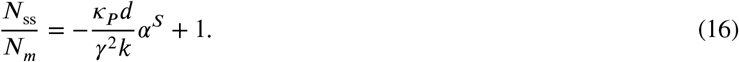

*Proof*. The steady-state value *N*_ss_ can be obtained more easily by considering the nondimensional model of (7)-(12) (see Appendix B for the complete details), setting the time derivatives to zero and solving the resulting algebraic equations. Such equations have only one admissible solution, the others being either the trivial solution or negative. The unique positive solution can be computed to be:

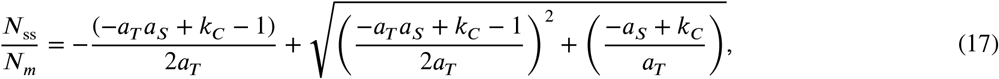

where

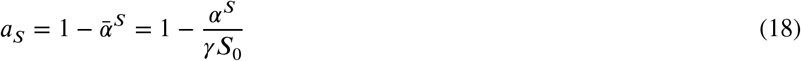

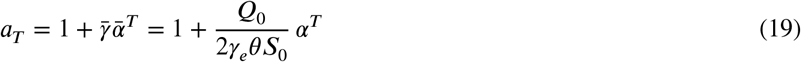

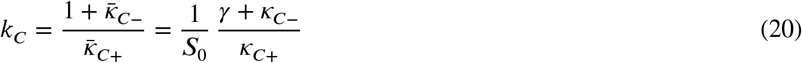

with *S*_0_ = (*γ k*) (*κ*_*P*_ *d*) and *Q*_0_ = (*N*_*m*_*ρ*_0_)/*γ* being nondimensional parameters (see Appendix B). Now, if the system’s parameters satisfy the following conditions

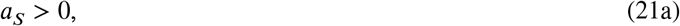

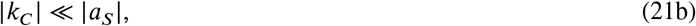

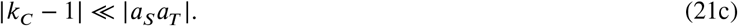

equation (17) can be recast as

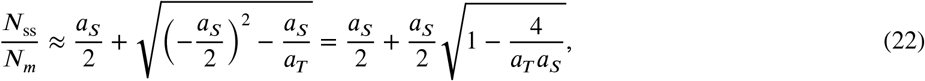

In addition to this, if

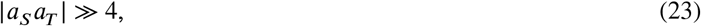

equation (22) can be further simplified yielding:

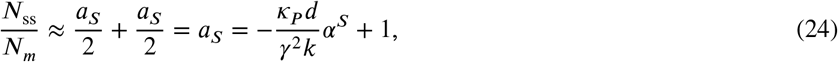

Indeed, after some algebraic manipulations, inequality (21a) can be rewritten as

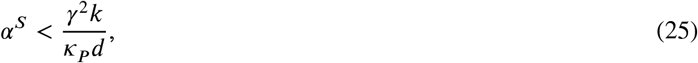

while (21b) as

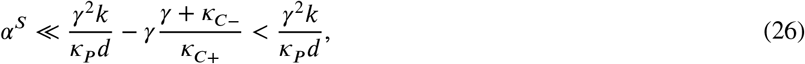

since *k*_*C*_ *>* 0. Therefore, under assumption (13), both the previous conditions (21a) and (21b) are verified.

Now, notice that assumption (14) implies that *k*_*C*_ *<* 1, and so also that 0 *<* | *k*_*C*_ − 1| *<* 1. Therefore, conditions (21c) and (23) can be rewritten as

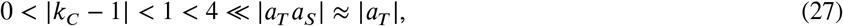

and thus both conditions (21c) and (23) are verified if

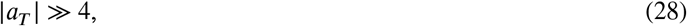

where the last step on the right hand-side in (27) follows from the fact that condition (26) implies that *a*_*S*_ ≈ 1. Substituting (19) into the last condition we obtain

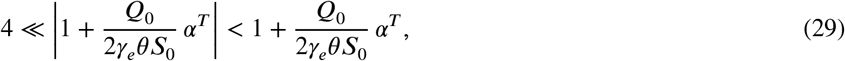

which is satisfied by assumption (15), thus completing the proof.

□

Note that assumptions (13) and (15) are equivalent to say, respectively, that the value of the reference parameter *α*^*S*^ must be small enough and that the production of the complex induced by the quorum sensing molecule is high enough. Additionally, condition (14) entails a timescale separation between the dynamics of the complex formation and of the population growth.

A more accurate relationship linking *N*_ss_ and *α*^*S*^ can be obtained by means of a similar analysis as done in Proposition 1, in the case that the basal production rates 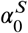 and 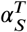 are non-zero, that is,

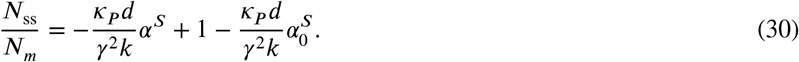

From Figure 2 it can be observed that equation (30) well approximates the input-output relationship at steady state between *N*_ss_/*N*_*m*_ and *α*^*S*^ of model (7)-(12). Specifically, provided that conditions (14) and (15) hold, the two curves overlap for values of *α*^*S*^ satisfying condition (13), which requires that *α*^*S*^ ≪ 191 trans h, for the values of the parameters we considered in this work (see Table A1).

**FIGURE 2.**
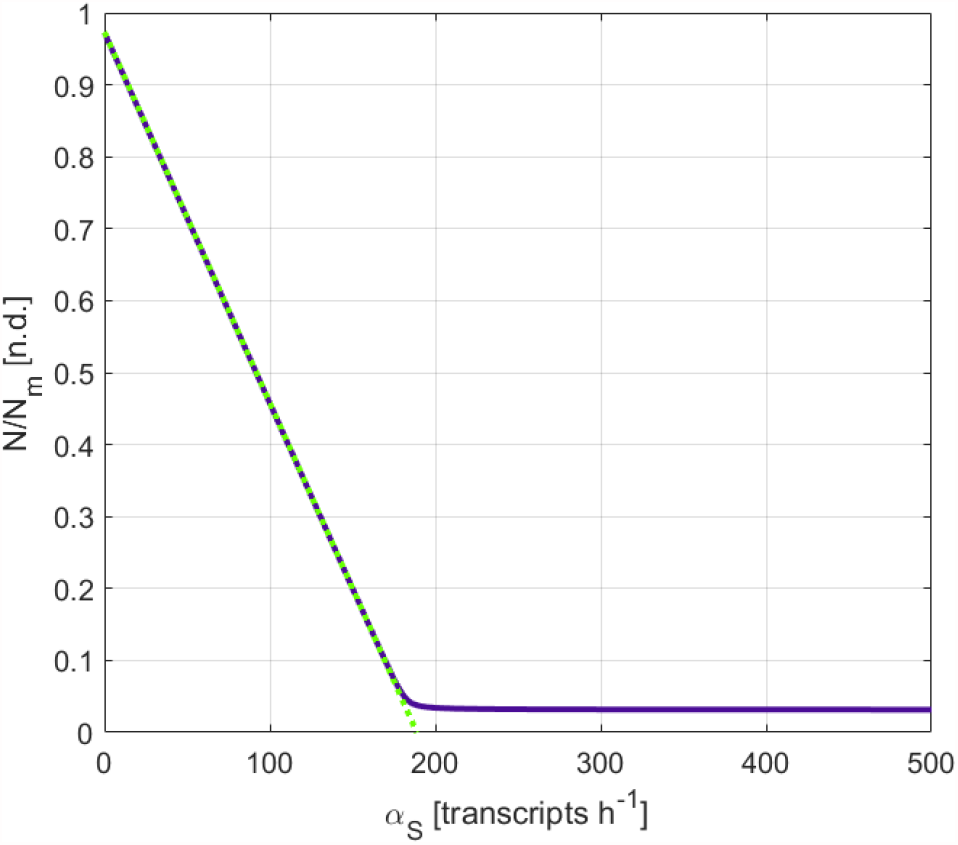
Steady-state input-output relationship between the normalized cellular density *N*_ss_/*N*_*m*_ and the reference parameter *α*^*S*^ obtained for the aggregated model (7)-(12) (blue line) and using equation (30) (dotted green line). Notice that the linear function in (30) gives a good approximation of the steady-state relationship for values of *α*^*S*^ satisfying (13), that is, *α*^*S*^ ≪ 191 trans h. The two curves were obtained by using the values of the parameters reported in Table A1.

By solving (30) for *α*^*S*^, we can obtain a useful expression to set the reference parameter *α*^*S*^ given the desired set-point of the population density at steady state, say *N*_d_, that is,

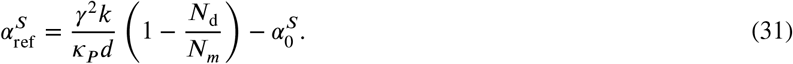

It can be noticed from the approximated expressions in (30) and (31) that the parameters appearing in the coefficient 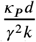 play a crucial role to achieve self-regulation to the desired set-point. In the next section, after validating the proposed embedded control architecture in different working conditions, we numerically investigate the robustness of our design with respect to parameters’ perturbations using model (7)-(12).

## 4 IN SILICO EXPERIMENTS

### 4.1 Agent-based simulations

To validate the effectiveness of the proposed population control strategy and to verify the accuracy of the analytical results obtained in Section 3, we implemented a set of *in silico* experiments by means of BSim ^28,29^, a realistic agent-based simulator of bacterial populations. BSim allows to simulate cells’ dynamics by also taking accounting for their spatial distribution and geometry, as well as for the spatio-temporal diffusion of signaling molecules into the environment and into the cells. In addition, it is also possible to simulate cell-to-cell variability, bio-mechanics, growth, division and death of the cells.

We considered a bacterial population, endowed with the proposed genetic regulatory networks encoding our control strategy, growing into an environment with limited availability of resources. Specifically, we assumed the culture environment to be a cube-shaped micro-vial with edges set to 10 *µ*m, that can at most host a population of 145 cells (Figure 3, right panel). This in order to strike a satisfactory trade-off between the computational cost of the simulations and the number of cells in the population. The numerical routine in BSim accounting for the cell growth dynamics was modified to replicate the typical logistic growth of cell populations under limited resources (further details are reported in Appendix C). The dynamics of each cell was implemented in BSim by using the agent-based model (1)-(5), complemented by the cell growth dynamics defined in the Appendix (see equations (C15) and (C19)). The nominal values of the parameters used in the simulations are reported in Table A1.

**FIGURE 3.**
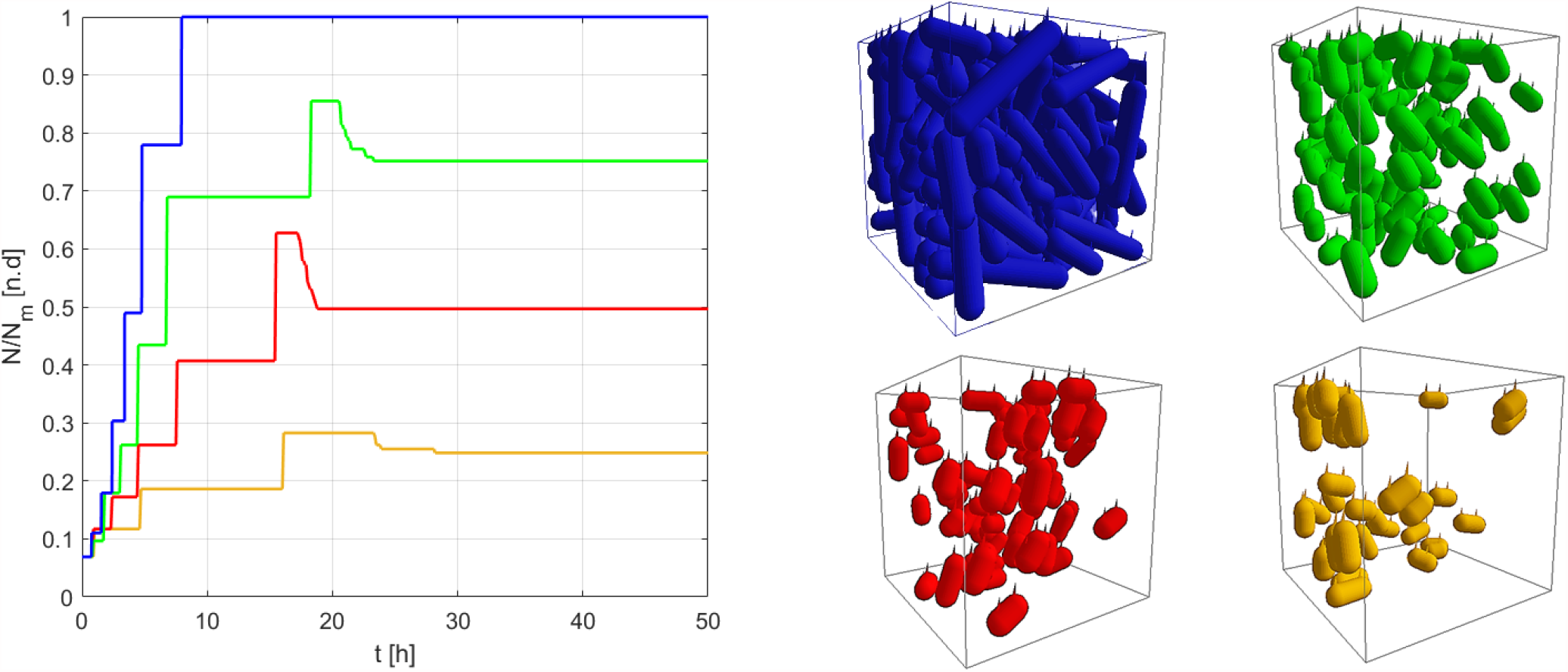
Set-point regulation experiments in BSim. *Left panel:* Time evolution of the population density in the cases of no growth control (blue line) and with growth control and desired set-point *N*_d_ equal to 0.25 *N*_*m*_ (yellow line), 0.50 *N*_*m*_ (red line), and 0.75 *N*_*m*_ (green line). The values for the desired density of the bacterial population were selected by choosing *α*^*s*^ as in Equation (31). *Right panel:* Snapshots of the culture chamber at steady state. All cells are assumed to be identical, that is, having all the parameters equal to their nominal values, as reported in Table A1. In all simulations we considered an initial population of 10 cells randomly arranged in the chamber.

### 4.2 Set-point regulation

We started by carrying out open-loop control experiments. As expected, in the absence of an embedded controller, the cell population density reaches the carrying capacity of the environment (Figure 3, left panel, blue line) with the cells filling completely the cubic chamber, that is, *N*_ss_ = *N*_*m*_ (Figure 3, right panel, top left chamber).

We then validated the proposed embedded genetic controller for three different set-points of the population density *N*_d_, namely 25%, 50% and 75% of the carrying capacity *N*_*m*_, assuming, to start with, all the cells to be *identical*, that is, with all parameters being equal to their nominal values (Table A1). The reference parameter 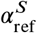, required to reach the desired set-point *N*_d_ / *N*_*m*_, was tuned according to the linear approximation in (31). In practice, this can be achieved *in vivo* either by engineering it via an offline design or by means of an additional external control loop (see for instance the external control of a tunable promoter proposed in 33. The *in silico* experiments confirmed that, under nominal conditions, the proposed control strategy allows the cells to self-regulate their density to the desired set-points by dynamically adapting their growth rate (Figure 3). Moreover, as shown in Figure 4, the control system was found to be robust to both sudden addition or removal of cells from the chamber; being able to recover the desired set-point after a short transient. Next we test its robustness to parameter variations.

**FIGURE 4.**
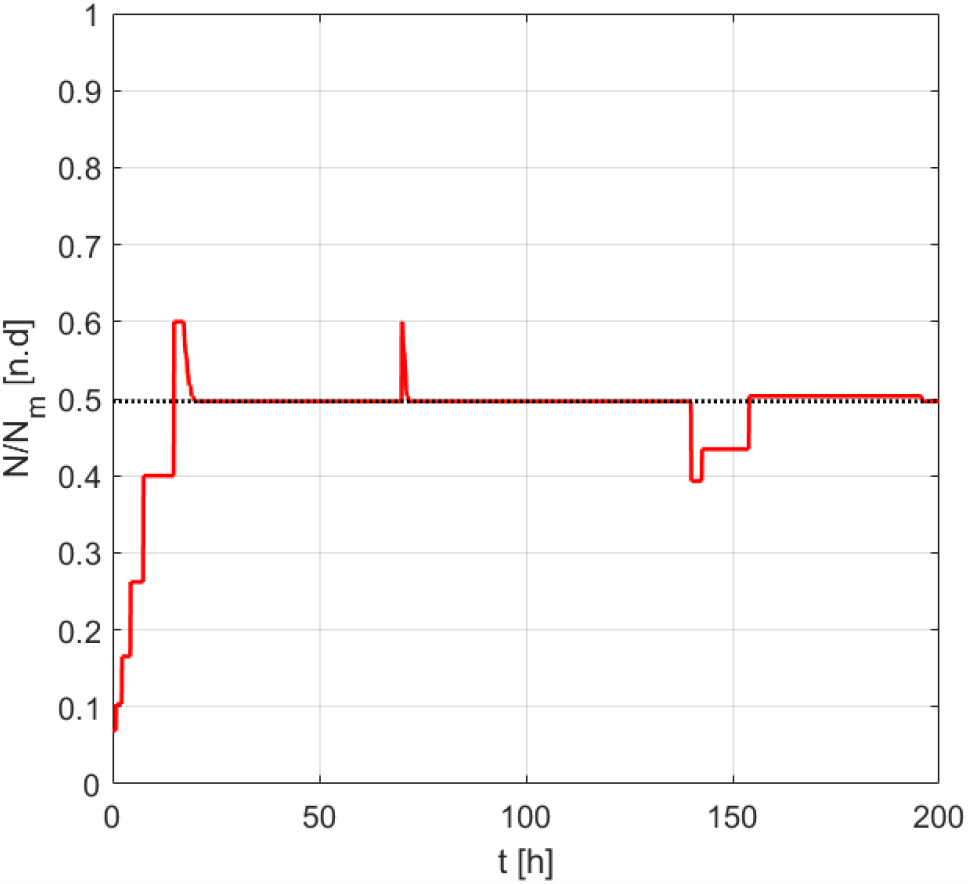
Robustness to impulsive disturbances. Time evolution of the population density of the closed-loop agent-based system in BSim in the presence of addition (at *t* = 70 h) and removal (at *t* = 140 h) of 15 cells (corresponding to about 0.1 *N*_*m*_).

### 4.3 Robustness to parameter variations

We analyzed the sensitivity of the controlled cells to variations of the parameters from their nominal values by running a series of simulations in MATLAB on the aggregate model (7)-(12). Specifically, we set the desired set-point to 50% of the carrying capacity, that is, *N*_d_ = 0.5 *N*_*m*_. Then, for any given parameter, say *µ*, and for each value of the coefficient of variation *CV*, we ran 200 simulations by drawing each time a parameter value from a normal distribution centered on its nominal value 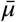 and with standard deviation 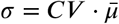, while keeping all the other parameters fixed to their nominal values. For each value of the coefficient of variation *CV* we considered, we evaluated the average percentage error at steady state over all the 200 simulations:

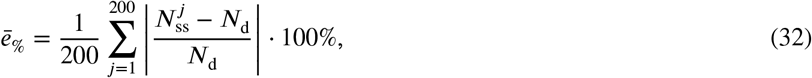

where 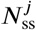 is the density at steady state reached by the cell population in the *j*-th simulation trial. The result of this analysis for *CV* = {0.05, 0.1, 0.15, 0.2} is reported in Figure 5 showing that only a subset of parameters significantly affects the steady-state error with a maximal variation relative to the unperturbed case of up to about 16% when *CV* = 0.2 for all parameters but *γ* which causes the largest variation (up to 30.44 % for *CV* = 0.2).

**FIGURE 5.**
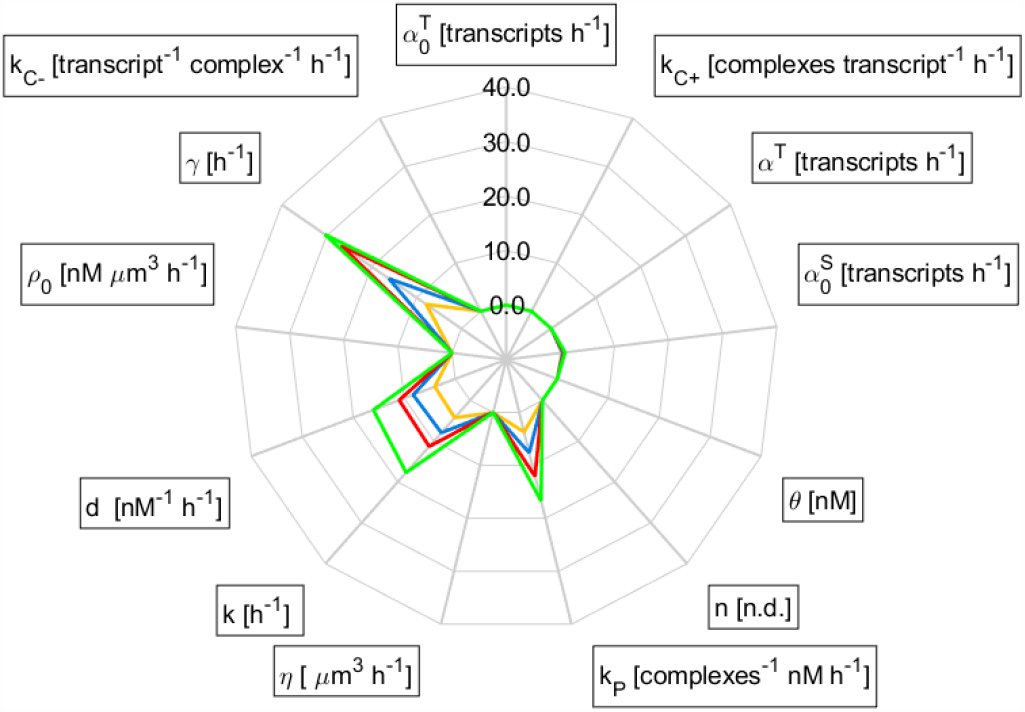
Robustness to parameter variations. Average percentage error at steady state *ē*_%_ of the aggregated model (7)-(12) as a function of the variability *CV* of the parameters’ value, for *CV* equal to 0.05 (yellow line), 0.1 (blue line), 0.15 (red line) and 0.2 (green line). For each parameter, say *µ*, and each value of *CV*, the results of 200 simulations were averaged, each obtained by drawing the value of *µ* from a normal distribution centered on its nominal value 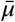 and with standard deviation 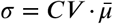, while keeping all the other parameters fixed to their nominal values. All simulations were run in MATLAB by setting the set-point *N*_d_ = 0.5 *N*_*m*_, initial density 0.1 *N*_*m*_, and on the time interval [0, 80 h].

Namely, these are the same parameters appearing in the linear approximations (16) and (30), that is, the growth rate *k*, the death rate *d*, the rate of production of the growth inhibitor protein *κ*_*P*_, and the dilution rate *γ*.

### 4.4 Cell-to-cell variability

Finally, cell-to-cell variability was modeled in BSim by assigning a different set of values of the parameters to daughter cells when they split from their mothers. We assumed that the heterogeneity in the response of the cells is essentially due to (i) the different copy numbers of the artificial plasmids into the cells and (ii) the different effects that the growth inhibitor protein *P* can have on the growth rate of each cell. Therefore, the parameters we considered to be varied in the agent-based model (1)-(5), and growth dynamics defined by (C15) and (C19), were 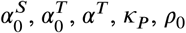, and the death rate *d* due to *P*. Specifically, each of these parameters, say *µ*, was drawn independently from a normal distribution centered on its nominal value 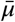 and with standard deviation 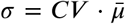. From the numerical experiments we observed that the average percentage error ē_%_, obtained over 100 simulation trials for each value of the set-point *N*_d_ and coefficient of variation *CV*, increases with the heterogeneity among the cells with lower values of the set-point *N*_d_ corresponding to higher sensitivity to cell-to-cell variability (Figure 6). Hence, the proposed controller can guarantee acceptable steady state values of the error in perturbed conditions when the reference set-point is greater than 40% while exhibiting larger deviations from the unperturbed case when the desired density goes below that value. In all cases, at the population level the cells exhibited very low variance over 100 trials, represented in Figure 6 by very small whiskers at each point.

**FIGURE 6.**
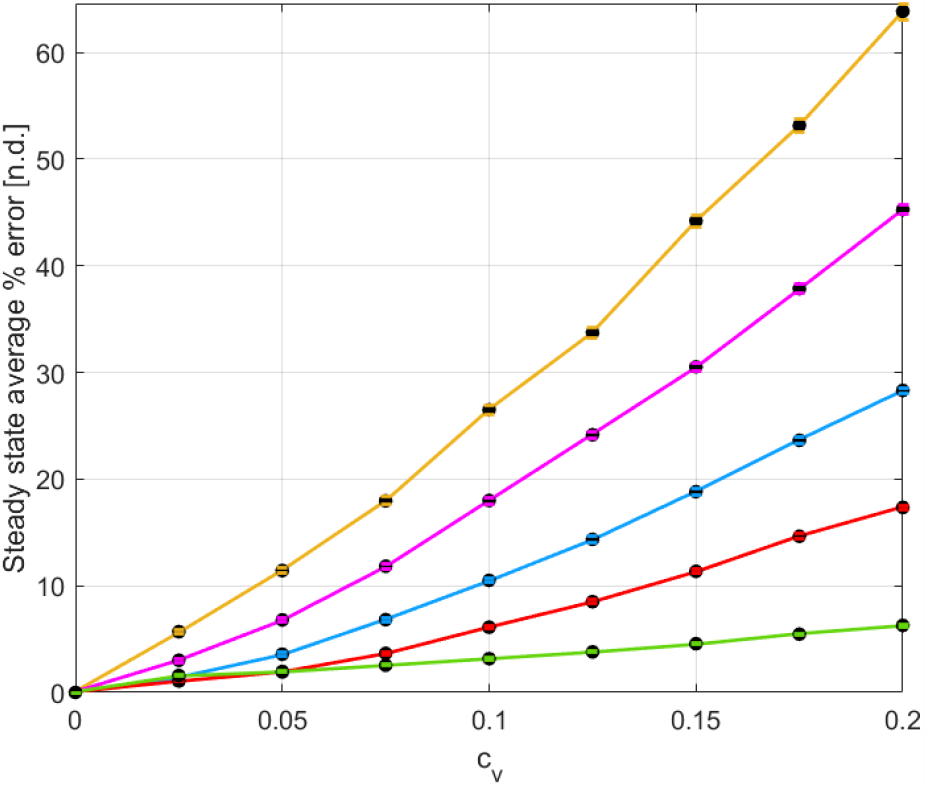
Cell-to-cell variability. Average value and standard deviation of the percentage error at steady state *-*_%_ of a heterogeneous population of cells (1)-(5), for the desired set-point *N*_d_ equal to 0.25 *N*_*m*_ (yellow), 0.30 *N*_*m*_ (pink), 0.40 *N*_*m*_ (blue), 0.50 *N*_*m*_ (red), and 0.75 *N*_*m*_ (green). Each point and its whiskers represent the average value and the standard deviation of ē_%_ evaluated over 100 simulation trials in BSim. For each simulation, all cells’ parameters were drawn independently from normal distributions centered on their nominal values, 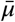, and with standard deviation 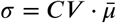. All simulations were initialized with a population of 7 cells. and the initial value of the state variables in (1)-(5) set to zero.

## 5 CONCLUSIONS

We considered the problem of regulating the growth rate of a bacterial population to reach a desired density at steady state. By engineering a tunable synthetic gene pathway, we were able to endow a cell population with the ability of self-regulating its own density to some desired value. We used a combination of analytical derivations and *in silico* experiments to test the performance and robustness of the proposed control architecture showing its reliability and robustness to parameter perturbations and cell-to-cell variability.

The genetic embedded controller presented in this paper was able to steer the population density to an arbitrary desired value, granting adaptation to loss or addition of cells while keeping the density close to the desired value even in the presence of cell-to-cell variability.

The proposed embedded strategy can therefore provide a solution for the self-regulation of cellular density that avoids unwanted extinction/overgrowth events. Our design impinges on the use of a unique tunable expression system that, contrary to other existing designs, exploits a toehold switch mechanism to allow simultaneous control of transcription and translation. Here this system is further engineered to provide a solution to self-regulation of cell growth that was shown to be effective and robust.

However, providing the correct reference to reach a desired set-point may be challenging in the presence of a heterogeneous cell population. To mitigate this problem, a promising solution may be to add an additional external feedback control loop which compensates for the heterogeneity by adjusting the reference fed to the inner system. This can be implemented *in vivo* by using the external control strategies ^34–39^.

The growth control approach we presented can be used either to substitute external feedback control loops to guarantee growth regulations of cell populations in chemostats or to robustify existing external feedback control strategies, providing some regulation capability in the presence of critical failures of the sensing or actuation systems. Another possible application is for ensuring long-term coexistence of different cell populations in microbial consortia for multicellular control (e.g., see 40,41). As such, the embedded control strategy we proposed can be used as an alternative providing a good trade-off between strategies allowing an online tuning of the consortium composition by modifying the external environment ^42^ and solutions which grant self-regulation of the consortium composition without allowing for the consortium composition to be changed dynamically ^25^.

□

## APPENDIX

### A DERIVATION OF THE AGGREGATE MODEL

The first part of the model, equations (7)-(10), follows immediately from the assumption that the average concentrations follow the same dynamics described in (1)-(4) for all cells in the population.

Equation (11) can be obtained by noticing that (5) can be rewritten as

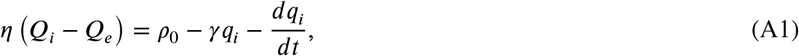

that substituted into (6) gives

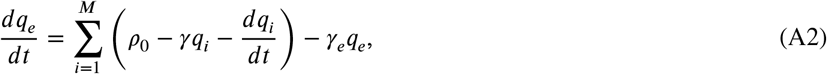

which in turns can be rewritten as

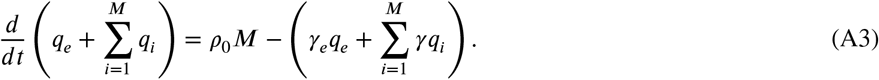

Now, noting that, by definition, 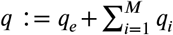, is the total amount of molecule Q into the external environment and into all the cells, and that, when considering a constant volume *V*, only degradation must be considered, that is, *γ* = *γ*_*e*_, we obtain

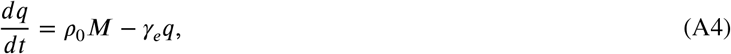

that, by dividing both sides by *V* and by using *N* = *M*/*V*, yields (11). Finally, equation (12) follows by assuming the typical logistic growth dynamics for the cell population in a limited environment, with the term −*dP N* capturing the slowdown effect of the inhibitor protein P on the growth rate of the cells, similarly to what was reported in 18.

**TABLE A1.** Nominal vaules of the systems’ parameters.

| Parameter | Value | Unit | Reference |
| --- | --- | --- | --- |
| $\alpha_0^S$ | 5.316 | tr h<sup>-1</sup> | 27 |
| $\alpha^S$ | variable | tr h<sup>-1</sup> | - |
| $\alpha_0^T$ | 13.4 | tr h<sup>-1</sup> | 27 |
| $\alpha^T$ | 11400 | tr h<sup>-1</sup> | 27 |
| $\kappa_{C+}$ | 1.542 | co tr<sup>-1</sup> h<sup>-1</sup> | 27 |
| $\kappa_{C-}$ | 0.4032 | co<sup>-1</sup> tr h<sup>-1</sup> | 27 |
| $\kappa_P$ | 5 | nM co<sup>-1</sup> h<sup>-1</sup> | 18 |
| $\rho_0$ | 595.2 | nM $\mu\text{m}^3$ h<sup>-1</sup> | - |
| $\gamma$ | 2 | h<sup>-1</sup> | 18 |
| $\gamma_e$ | 2 | h<sup>-1</sup> | 18 |
| $\eta$ | 120 | $\mu\text{m}^3$ h<sup>-1</sup> | - |
| $\theta$ | 20 | nM | - |
| $n$ | 2 | - | 27 |
| $k$ | 0.97 | h<sup>-1</sup> | 18 |
| $d$ | $4 \cdot 10^{-3}$ | nM<sup>-1</sup> h<sup>-1</sup> | 18 |
| $N_m$ | 0.145 | CFU $\mu\text{m}^{-3}$ | - |
| $V_c$ | 5.5 | $\mu\text{m}^3$ | - |

The units of measurement tr, co and CFU stand for transcripts, complexes, and colony-forming units, respectively.

### B NONDIMENSIONAL MODEL

Model (7)-(12) can be nondimensionalized to obtain a simpler model with less parameters. Specifically, by assuming that the Hill function in (2) is not saturated, i.e., *Q* ≈ *θ*, the activation function of *T* due to *Q* in (2) can be approximated by a linear function, that is,

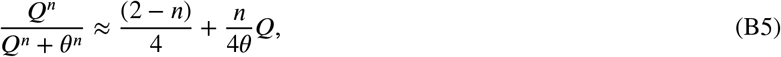

which reduceds to 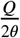 for *n* = 2. Then, rescaling time by setting *τ* = *γt* and introducing the following dimesionless variables:

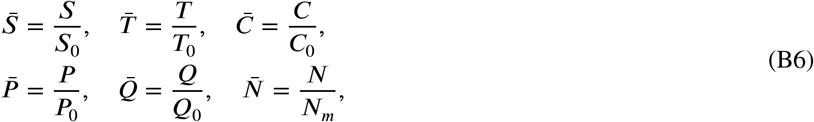

with

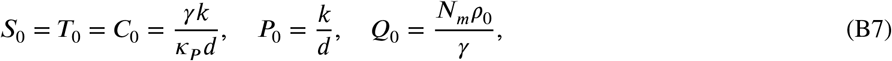

a *nondimensional model* of (7)-(12) can be obtained as:

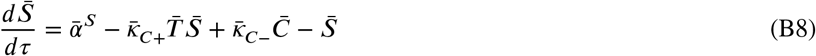

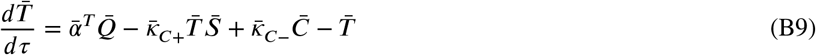

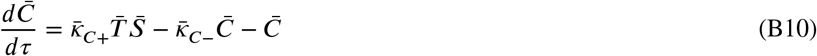

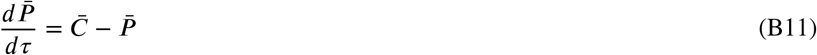

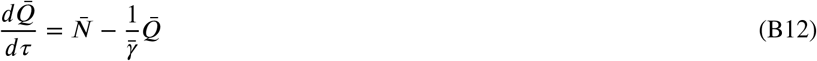

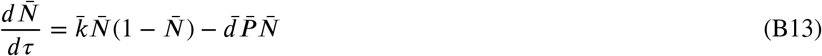

where

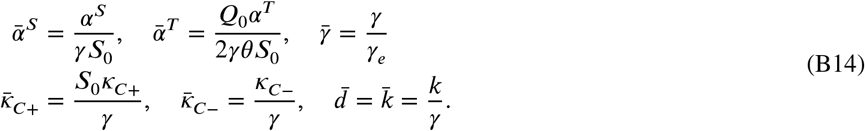

are positive nondimensional parameters.

### C LOGISTIC GROWTH DYNAMICS IN BSIM

The typical rod-shaped geometry of *E. coli* bacteria is modeled in BSim by considering for each cell a cylinder with radius *r* = 0.5 *µ*m and length *L* having a semi-sphere with the same radius *r* attached to both bases. Cells have a random initial length drawn with uniform distribution from the interval [0.7, 1.3] *µ*m, and their growth is implemented by assuming that the length *L* of each cell grows by following a logistic dynamics of the form:

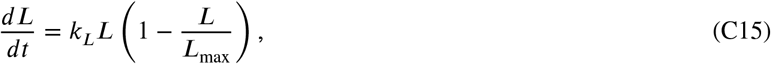

where *L*_max_ denotes the maximum cell length, set to 5.625 *µ*m, and *k*_*L*_ denotes the growth rate. Cell division occurs when the length of the cell reaches the critical threshold ^29, SI^ *L*_th_ = 4.5 *µ*m. As a consequence, the time needed for the division to occur, that is, the doubling time of the population, given the initial length of a bacterium *L*_0_, is defined as:

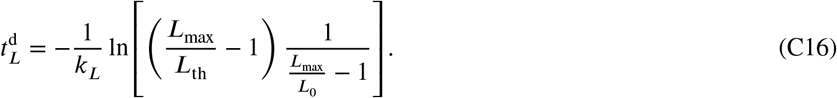

Note that, in the original implementation in BSim, 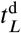 does not depend on the population density *N*, hence population will grow exponentially at a constant rate. Therefore, in order for the population density *N* to follow a logistic growth rate in BSim, it is necessary to make *k*_*L*_ depend directly upon *N*. To this aim, we assume that the doubling time of the cell length 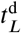 is equal to the doubling time of the population density, whose dynamics is described by:

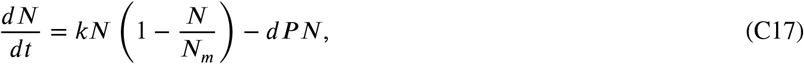

so that the doubling time of the population density can be derived as:

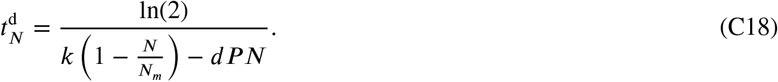

Finally, by equating the two doubling times, 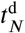 and 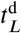, we obtain the desired relationship between *k*_*L*_ and *N* as:

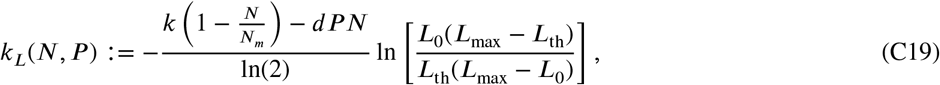

which was implemented to carry out the agent-based simulations reported to the research.

## ACKNOWLEDGMENTS

Mario di Bernardo, Davide Fiore and Davide Salzano wish to acknowledge support from the European Union’s Horizon 2020 research and innovation programme under grant agreement No 766840 (COSY-BIO). Davide Fiore also wishes to acknowledge support by the research grant “BIOMASS” from the University of Naples Federico II - “Finanziamento della Ricerca di Ateneo (FRA) - Linea B”.

## Author contributions

Virginia Fusco and Davide Salzano contributed equally to this research.

## 0 Abbreviations

TES: tunable expression system

## References

1. Brenner K, You L, Arnold FH. Engineering microbial consortia: a new frontier in synthetic biology. Trends in Biotechnology 2008; 26(9): 483–489. doi: 10.1016/j.tibtech.2008.05.004

2. Satyanarayana T, Kunze G. Yeast biotechnology: diversity and applications. Springer, Dordrecht . 2009.

3. Su Y, Liu C, Fang H, Zhang D. Bacillus subtilis: a universal cell factory for industry, agriculture, biomaterials and medicine. Microbial Cell Factories 2020; 19(173). doi: 10.1186/s12934-020-01436-8

4. Jullesson D, David F, Pfleger B, Nielsen J. Impact of synthetic biology and metabolic engineering on industrial production of fine chemicals. Biotechnology Advances 2015; 33(7): 1395–1402. doi: 10.1016/j.biotechadv.2015.02.011

5. Choi Y, Park TJ, Lee DC, Lee SY. Recombinant Escherichia coli as a biofactory for various single-and multi-element nano-materials. Proceedings of the National Academy of Sciences 2018; 115(23): 5944–5949. doi: 10.1073/pnas.1804543115

6. Hug JJ, Krug D, Müller R. Bacteria as genetically programmable producers of bioactive natural products. Nature Reviews Chemistry 2020; 4: 172–193. doi: 10.1038/s41570-020-0176-1

7. Mauri M, Gouzé JL, De Jong H, Cinquemani E. Enhanced production of heterologous proteins by a synthetic microbial community: Conditions and trade-offs. PLoS Comput Biol 2020; 16(4): e1007795. doi: 10.1371/journal.pcbi.1007795

8. Lo TM, Chng SH, Teo WS, Cho HS, Chang MW. A two-layer gene circuit for decoupling cell growth from metabolite production. Cell Systems 2016; 3(2): 133–143. doi: 10.1016/j.cels.2016.07.012

9. Tian R, Liu Y, Cao Y, et al. Titrating bacterial growth and chemical biosynthesis for efficient N-acetylglucosamine and N-acetylneuraminic acid bioproduction. Nature Communications 2020; 11(5078). doi: 10.1038/s41467-020-18960-1

10. Xu P. Production of chemicals using dynamic control of metabolic fluxes. Current Opinion in Biotechnology 2018; 53: 12–19. doi: 10.1016/j.copbio.2017.10.009

11. Lv Y, Qian S, Du G, Chen J, Zhou J, Xu P. Coupling feedback genetic circuits with growth phenotype for dynamic population control and intelligent bioproduction. Metabolic Engineering 2019; 54: 109–116. doi: 10.1016/j.ymben.2019.03.009

12. Monod J. The growth of bacterial cultures. Annual Review of Microbiology 1949; 3(1): 371–394. doi: 10.1146/annurev.mi.03.100149.002103

13. De Leenheer P, Smith H. Feedback control for chemostat models. Journal of Mathematical Biology 2003; 46(1): 48–70. doi: 10.1007/s00285-002-0170-x

14. Fiore D, Della Rossa F, Guarino A, di Bernardo M. Feedback ratiometric control of two microbial populations in a single chemostat. IEEE Control Systems Letters 2021: 1–1. doi: 10.1109/LCSYS.2021.3086234

15. Zhu GY, Zamamiri A, Henson MA, Hjortsø MA. Model predictive control of continuous yeast bioreactors using cell population balance models. Chemical Engineering Science 2000; 55(24): 6155–6167. doi: 10.1016/S0009-2509(00)00208-6

16. Richmond A, Cheng-Wu Z, Zarmi Y. Efficient use of strong light for high photosynthetic productivity: interrelationships between the optical path, the optimal population density and cell-growth inhibition. Biomolecular Engineering 2003; 20(4-6): 229–236. doi: 10.1016/S1389-0344(03)00060-1

17. Treloar NJ, Fedorec AJ, Ingalls B, Barnes CP. Deep reinforcement learning for the control of microbial co-cultures in bioreactors. PLoS Comput Biol 2020; 16(4): e1007783. doi: 10.1371/journal.pcbi.1007783

18. You L, Cox RS, Weiss R, Arnold FH. Programmed population control by cell–cell communication and regulated killing. Nature 2004; 428(6985): 868–871. doi: 10.1038/nature02491

19. Fedorec AJ, Karkaria BD, Sulu M, Barnes CP. Single strain control of microbial consortia. Nature Communications 2021; 12(1977). doi: 10.1038/s41467-021-22240-x

20. Alnahhas RN, Sadeghpour M, Chen Y, et al. Majority sensing in synthetic microbial consortia. Nature Communications 2020; 11(3659). doi: 10.1038/s41467-020-17475-z

21. Stephens K, Pozo M, Tsao CY, Hauk P, Bentley WE. Bacterial co-culture with cell signaling translator and growth controller modules for autonomously regulated culture composition. Nature Communications 2019; 10(4129). doi: 10.1038/s41467-019-12027-6

22. Karkaria BD, Fedorec AJ, Barnes CP. Automated design of synthetic microbial communities. Nature Communications 2021; 12(672). doi: 10.1038/s41467-020-20756-2

23. Dinh CV, Chen X, Prather KL. Development of a quorum-sensing based circuit for control of coculture population composition in a naringenin production system. ACS Synthetic Biology 2020; 9(3): 590–597. doi: 10.1021/acssynbio.9b00451

24. Scott SR, Din MO, Bittihn P, Xiong L, Tsimring LS, Hasty J. A stabilized microbial ecosystem of self-limiting bacteria using synthetic quorum-regulated lysis. Nature Microbiology 2017; 2(17083). doi: 10.1038/nmicrobiol.2017.83

25. Ren X, Baetica AA, Swaminathan A, Murray RM. Population regulation in microbial consortia using dual feedback control. In: Proc. of the 56th IEEE Conference on Decision and Control. ; 2017: 5341–5347. doi: 10.1109/CDC.2017.8264450

26. McCardell RD, Pandey A, Murray RM. Control of density and composition in an engineered two-member bacterial community. bioRxiv 2019: 632174. doi: 10.1101/632174

27. Bartoli V, Meaker GA, di Bernardo M, Gorochowski TE. Tunable genetic devices through simultaneous control of transcription and translation. Nature Communications 2020; 11(2095). doi: 10.1038/s41467-020-15653-7

28. Gorochowski TE, Matyjaszkiewicz A, Todd T, et al. BSim: an agent-based tool for modeling bacterial populations in systems and synthetic biology. PloS One 2012; 7(8): e42790. doi: 10.1371/journal.pone.0042790

29. Matyjaszkiewicz A, Fiore G, Annunziata F, et al. BSim 2.0: an advanced agent-based cell simulator. ACS Synthetic Biology 2017; 6(10): 1969–1972. doi: 10.1021/acssynbio.7b00121

30. Chen Y, Ho JM, Shis DL, et al. Tuning the dynamic range of bacterial promoters regulated by ligand-inducible transcription factors. Nature Communications 2018; 9(64). doi: 10.1038/s41467-017-02473-5

31. Hughes RM. A compendium of chemical and genetic approaches to light-regulated gene transcription. Critical Reviews in Biochemistry and Molecular Biology 2018; 53(5): 453–474. doi: 10.1080/10409238.2018.1487382

32. Baumschlager A, Rullan M, Khammash M. Exploiting natural chemical photosensitivity of anhydrotetracycline and tetracycline for dynamic and setpoint chemo-optogenetic control. Nature Communications 2020; 11(3834). doi: 10.1038/s41467-020-17677-5

33. Menolascina F, Fiore G, Orabona E, et al. In-vivo real-time control of protein expression from endogenous and synthetic gene networks. PLoS Comput Biol 2014; 10(5): e1003625. doi: 10.1371/journal.pcbi.1003625

34. Scott TD, Sweeney K, McClean MN. Biological signal generators: integrating synthetic biology tools and in silico control. Current Opinion in Systems Biology 2019; 14: 58–65. doi: 10.1016/j.coisb.2019.02.007

35. Chen S, Harrigan P, Heineike B, Stewart-Ornstein J, El-Samad H. Building robust functionality in synthetic circuits using engineered feedback regulation. Current Opinion in Biotechnology 2013; 24(4): 790–796. doi: 10.1016/j.copbio.2013.02.025

36. Milias-Argeitis A, Rullan M, Aoki SK, Buchmann P, Khammash M. Automated optogenetic feedback control for precise and robust regulation of gene expression and cell growth. Nature Communications 2016; 7(12546). doi: 10.1038/ncomms12546

37. Shannon B, Zamora-Chimal CG, Postiglione L, et al. In vivo Feedback Control of an Antithetic Molecular-Titration Motif in Escherichia coli using Microfluidics. ACS Synthetic Biology 2020; 9(10): 2617–2624. doi: 10.1021/acssynbio.0c00105

38. Guarino A, Fiore D, Salzano D, di Bernardo M. Balancing cell populations endowed with a synthetic toggle switch via adaptive pulsatile feedback control. ACS Synthetic Biology 2020; 9(4): 793–803. doi: 10.1021/acssynbio.9b00464

39. Harrigan P, Madhani HD, El-Samad H. Real-time genetic compensation defines the dynamic demands of feedback control. Cell 2018; 175(3): 877-886.e10. doi: 10.1016/j.cell.2018.09.044

40. Fiore D, Salzano D, Cristòbal-Cóppulo E, Olm JM, di Bernardo M. Multicellular feedback control of a genetic toggle-switch in microbial consortia. IEEE Control Systems Letters 2020; 5(1): 151–156. doi: 10.1109/LCSYS.2020.3000954

41. Fiore G, Matyjaszkiewicz A, Annunziata F, et al. In-silico analysis and implementation of a multicellular feedback control strategy in a synthetic bacterial consortium. ACS Synthetic Biology 2017; 6(3): 507–517. doi: 10.1021/acssynbio.6b00220

42. Salzano D, Fiore D, di Bernardo M. Ratiometric control for differentiation of cell populations endowed with synthetic toggle switches. In: Proc. of 58th IEEE Conference on Decision and Control. ; 2019: 927–932. doi: 10.1109/CDC40024.2019.9029592

